# Secondary environmental variation creates a shifting evolutionary watershed for the methyl-parathion hydrolase enzyme

**DOI:** 10.1101/833764

**Authors:** Dave W. Anderson, Florian Baier, Gloria Yang, Nobuhiko Tokuriki

## Abstract

Enzymes can evolve new catalytic activity when their environments change to present them with novel substrates. Despite this seemingly straightforward relationship, factors other than the direct catalytic target can also impact enzyme adaptation. Here, we characterize the adaptive landscape separating an ancestral dihydrocoumarin hydrolase from a methyl parathion hydrolase descendant under eight different environments supplemented with alternative divalent metals. This variation shifts an evolutionary watershed, causing the outcome of adaptation to depend on the environment in which it occurs. The resultant landscapes also vary in terms both the number and the genotype(s) of “fitness peaks” as a result of genotype-by-environment (*G×E*) interactions and environment-dependent epistasis (*G×G×E*). This suggests that adaptive landscapes may be fluid and that molecular adaptation is highly contingent not only on obvious factors (such as catalytic targets) but also on less obvious secondary environmental factors that can direct it toward distinct outcomes.

## Introduction

Evolution is fundamentally dynamic, encompassing the myriad of ways in which biological forms and functions change from one state into another.^1–3^ Species or genes undergoing the classic Darwinian model of adaptation can be visualized as akin to rainwater in a drainage basin, which travels along the surface of the Earth until it flows into the ocean (Figure 1A).^4–6^ In the case of rainwater, the “drainage” is mostly predictable, even when rain initially falls hundreds of kilometers away from its ultimate destination; the land area that drains to the same destination is what we term a “watershed”. Under some circumstances, moving or re-shaping critical ridges on the landscape even slightly can shift these watersheds, causing rainwater falling on the same geographical position to drain into the Atlantic rather than the Pacific Ocean, for example.^7^ The concept of these watershed boundaries, and their potential to shift, are commonly applied in general to questions of causality and contingency, for example in history, where it is common to discuss so-called “watershed moments” (Figure 1A).^8^ If we draw an analogy to enzyme adaptation, we would consider each ocean to be the most fit genotype (*i.e.*, that which exhibits the optimal enzyme activity for its target substrate) available through the acquisition of function-altering mutations. In this analogy, “gravity” (the force causing rainwater to move across the geographical landscape) is akin to the selection pressure that is defined by the “primary” environment (*i.e.*, the novel substrate that is the target of a favourable catalytic function in that environment) (Figure 1B).^9,10^ But how fixed are these adaptive landscapes? Can they be re-shaped or altered by variation in “secondary” environmental factors, such as temperature, salinity, pH, the presence of other proteins, or cofactor availability such as metals? These factors do not necessarily define the novel adaptive function but they can nonetheless impact the fitness of a genotype;^11,12^ can they also impact the direction of their evolution (Figure 1B)?^13–15^ Several enzyme studies have addressed the impact of “primary” environments (*i.e.*, different substrates or ligands) on the topology of the adaptive landscapes,^16,17^ the degree to which the secondary environmental factors can alter evolutionary outcomes even under the same primary selective pressure remains poorly understood^11,18^). If natural selection drives the evolution of new forms or functions, under what circumstances can that outcome vary? Can secondary environmental variation shift key evolutionary watersheds, causing the same genetic starting point to evolve along different trajectories and, ultimately, to reach different adaptive outcomes?

**Figure 1:**
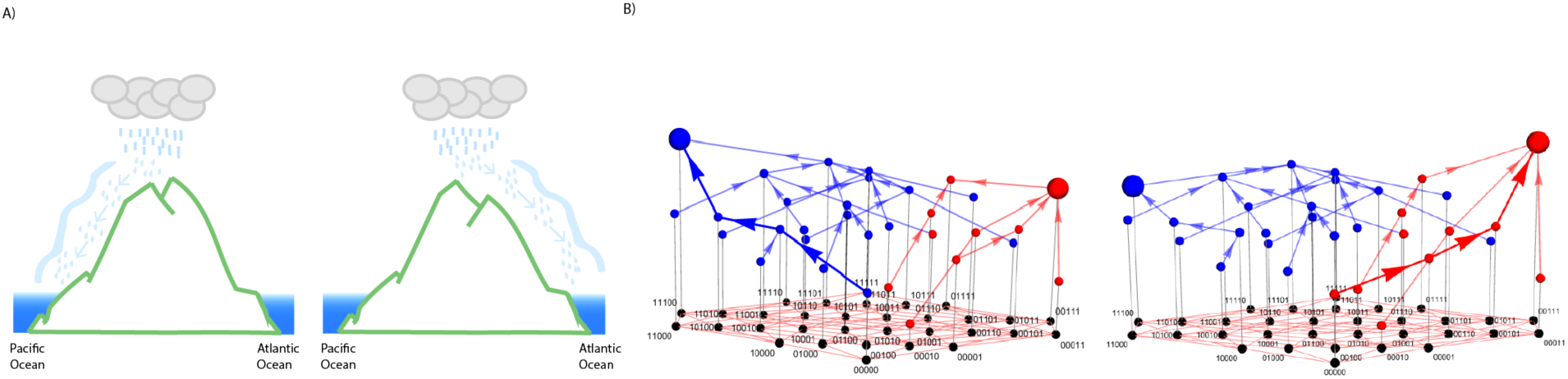
Schematic outline of analogy between geological watersheds (A) and evolutionary watersheds (B).

We explore these questions and concepts in detail by characterizing the evolutionary transition between an ancestral dihydrocoumarin hydrolase (DHCH) and its methyl-parathion hydrolase (MPH) descendant within the metallo-β-lactamase (MBL) superfamily^19^ under environmentally realistic secondary environments.^20^ This adaptation occurred between the 1940s and the 2000s, coinciding with the human application of organophosphate (OP) pesticides in industry and agriculture, thus providing an excellent case of classic Darwinian adaptation.^21,22^ Previous work characterized a set of five mutations – four single-nucleotide changes and one single-residue insertion that surround the active site (*l*72R, Δ193S, *h*258L, *i*271T, and *f*273L; Figure 2A) – that is both necessary and sufficient to recapitulate the evolution of the derived methyl-parathion hydrolase activity in a zinc-rich environment.^23^ Here, we investigate the impact of variation in a secondary environmental factor, specifically the type of metal ion present in the cellular environment, on adaptation from the DHCH ancestor. We systematically characterize the same genotypes (a complete combinatorial set of those five historical mutations – 32 genotypes in total) in order to further examine and compare the adaptive landscapes for each metal environment (Supplemental Figure 1). By combining this approach with diverse and extensive statistical analyses, we unveil the extent to which the secondary environment can shift evolutionary watersheds and potentially alter evolutionary trajectories and outcomes in this system.

**Figure 2:**
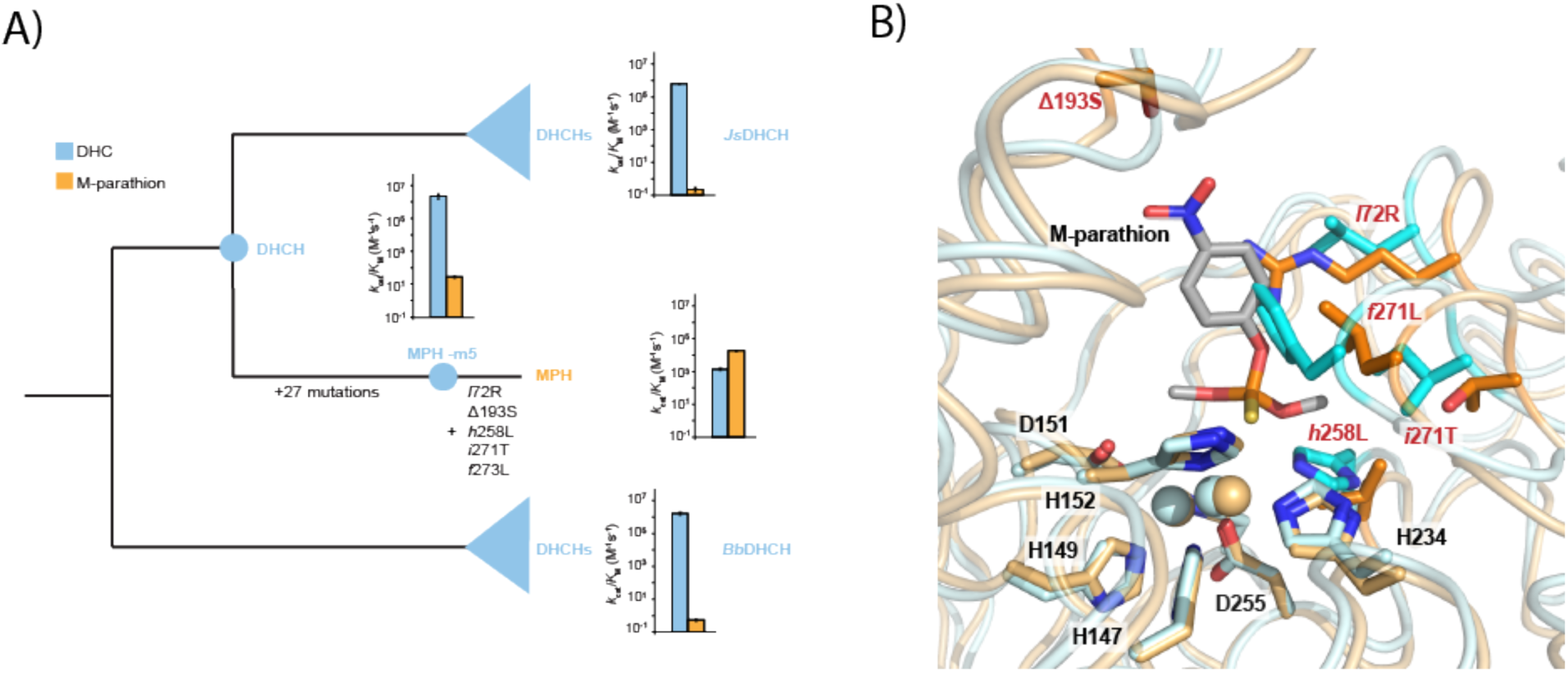
**A:** Phylogenetic reconstruction of MPH family and its DHCH relatives. Phylogenetic display was constructed using previously published larger phylogenetic reconstruction.^23^ **B:** The DHCH and MPH crystal structures aligned, with evolutionarily relevant substitutions highlighted.

## Results

### An experimental scheme reveals the effect of different metal environments on the evolution of MPH activity

We define the primary environment or direct selection pressure during MPH adaptation to be the maximized hydrolase activity against methyl-parathion. Both MPH and DHCH enzymes require two divalent ions to be coordinated in their active site in order to be catalytically active (Figure 2B).^23–25^ Whereas the majority of research on MPH to date assume it solely or primarily functions using Zn^2+^ to coordinate the substrate in its active site,^26^ MPH has also been shown to exhibit varying enzymatic activity and promiscuity when other divalent metals are also present.^24,25^ Thus, we define the secondary environment as the abundance of metal ions in the environment, and examine the effect of its variation on the adaptive landscape of MPH. We selected eight different environmentally-relevant divalent metals (calcium – Ca^2+^, cadmium – Cd^2+^, cobalt – Co^2+^, copper – Cu^2+^, magnesium – Mg^2+^, manganese – Mn^2+^, nickel – Ni^2+^, and zinc – Zn^2+^). It is known that these eight metal ions are commonly present in soil, particularly in industrial and agricultural environments where methyl-parathion is used and where MPH enzymes were originally discovered in soil bacteria,^20,27,28^ meaning our experiment reflects realistic alternative, secondary environments in which MPH adaptation could have occurred. We characterized the complete adaptive landscape defined by five key historical genetic changes – all 32 combinatorial genotypes that separate the ancestral DHCH created by taking the MPH genotype and reversing the five historical mutations) and the derived MPH under eight different secondary environments. All 32 genes were transformed and expressed in *E. coli* BL21 (DE3), which were grown in cell media supplemented with only one of eight divalent metals, and the methyl-parathion hydrolase (MPH) activity of cell lysate was measured by mixing with methyl-parathion and monitoring the appearance of the leaving group. Note that the LB media used for the cell culture contains some trace of these metal ions,^29^ and that adding additional supplemental metal (200 µM) would alter the proportion of metal ions in the cell while the metal concentrations are controlled, to a certain degree, by homeostasis mechanisms. Regardless, we have previously shown that supplementing media with divalent metals in this way does not affect the growth rate of *E. coli* but does impact the activity level of MPH variants.^24^ Still, it is likely that not all intracellular MPH enzymes are acquiring the supplemented metal in the cell (and in particular, some metal ions such as Ca^2+^ and Mg^2+^ may not associate strongly with the enzyme). Also, it is more likely that the enzymes are adopting a mixture of multiple metal-bound states, including each metal binding site accommodate different metal^24^ that may exhibit different catalytic activities. Nonetheless, what is clear, and what is most important for our study here, is that these metal environments significantly impact the catalytic activity level in the cell, and could therefore conceivably impact the process of enzyme adaptation.

### Variation in the adaptive landscape results in divergent adaptive outcomes

Each metal environment creates a unique adaptive landscape, and comparing them highlights several meaningful differences. First, different metal environments result in varying levels of methyl-parathion hydrolysis activity for the fully ancestral (DHCH) and descendant (MPH) enzymes, with more than 100-fold variation in methyl-parathion hydrolysis activity for DHCH and more than 10-fold variation for MPH (Figure 3A). Critically, the change in activity between DHCH and MPH also varies significantly, ranging from ~18-fold improvement (in the Ni^2+^ environment) to 910-fold improvement (in the Zn^2+^ environment), indicating that the effect of the five historical mutations varies significantly depending on the metal environment (Figure 3B). Second, the effect of even a single mutation in the ancestral genotypic background varies substantially depending on the metal environment (Figure 3C). For example, the effect of the *l*72R mutation is positive with seven metals, but negative with Mn^2+^. Similarly, *i*271T had a positive effect in the presence of Cd^2+^, but had a negative effect in all other metal environments. Moreover, whereas *h*258L has a consistently positive effect in all metal environments, the magnitude of its effect varies significantly ranging from ~18-fold improvement (in the Cu^2+^ environment) up to ~510-fold improvement (in the Mg^2+^ environment) (Figure 3C).

**Figure 3:**
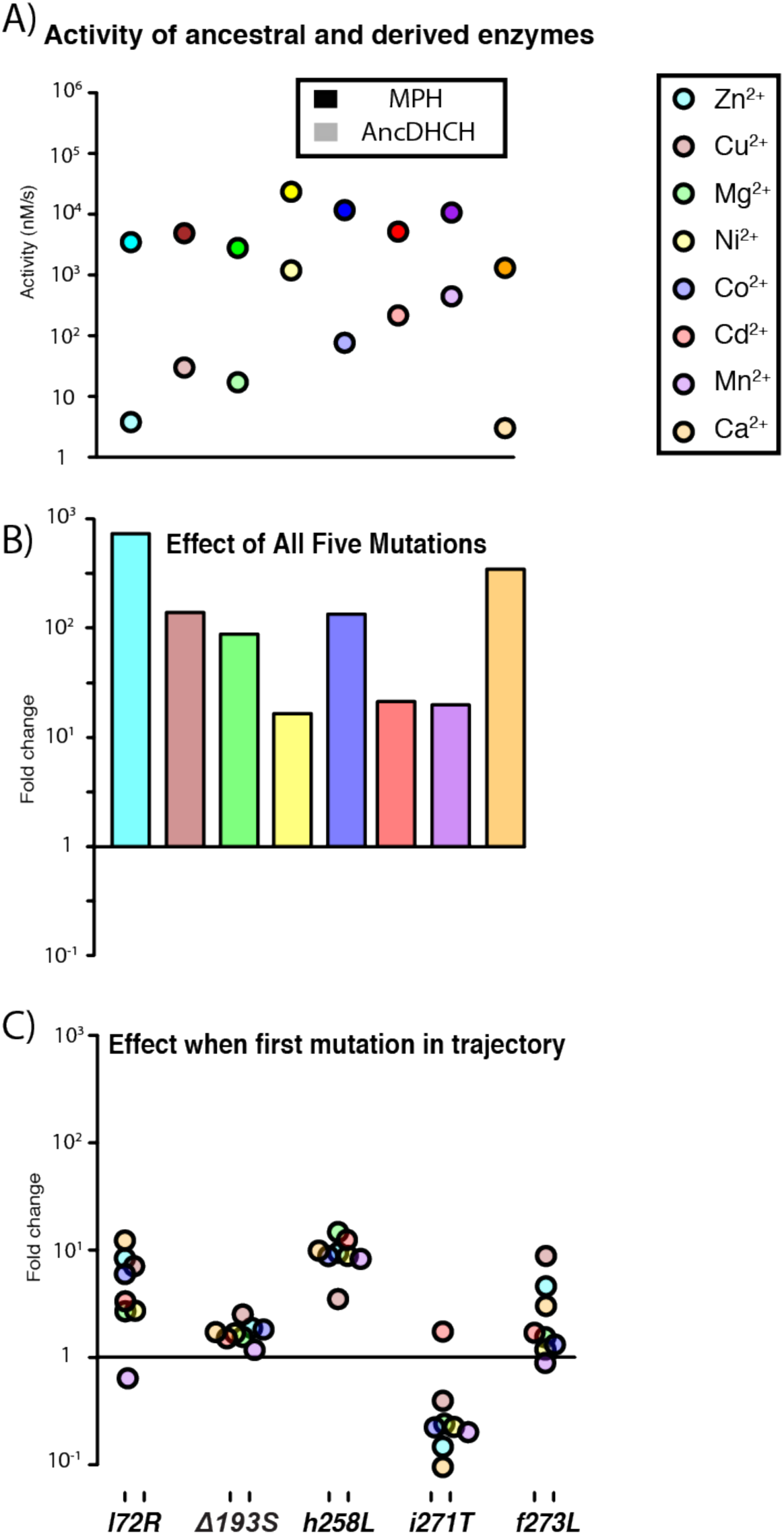
**A:** The catalytic activity for both the ancestral (faded circles) and derived (solid circles) enzymes when expressed in different metal environments. **B:** The collective effect of all five historical mutations in each metal environment. **C**: The effect of each individual mutation when introduced into the ancestral genetic background.

Moreover, the overall topology of the adaptive landscape in each metal environment differs substantially (Figure 4). To assess the consequences of this variation for the adaptive process, we applied a simple model of directional Darwinian selection to calculate the most likely trajectory beginning from the ancestral genotype across the adaptive landscape and ending at an “optimal” genotype (*i.e.*, from which all available single mutations would reduce MPH activity – see **Methods** and Figure 4).^30,31^ Interestingly, the evolution of MPH activity in different metal environments results in a trajectory that leads to different local optimal genotypes (Figure 4). For example, trajectories beginning at the ancestral genotype led to the fully derived MPH genotype in only four out of eight secondary environments (Ca^2+^, Co^2+^, Cu^2+^ and Zn^2+^ - Figure 4A,B,E,H). Of the remaining environments tested, three (Mg^2+^, Mn^2+^ and Ni^2+^ - Figure 4C,D,G) maintained the derived MPH as the global optimum across the landscape; however, for each of them the adaptive trajectory that begins from the ancestral DHCH genotype failed to reach it, instead becoming stranded on a local optimum. Finally, in one secondary environment (Cd^2+^) there was a unique global optimum that was not the fully derived MPH genotype (Figure 4F). Taken together, it is clear that variation in the metal environment can result in varying adaptive landscapes and, as a result, divergent evolutionary outcomes.

**Figure 4:**
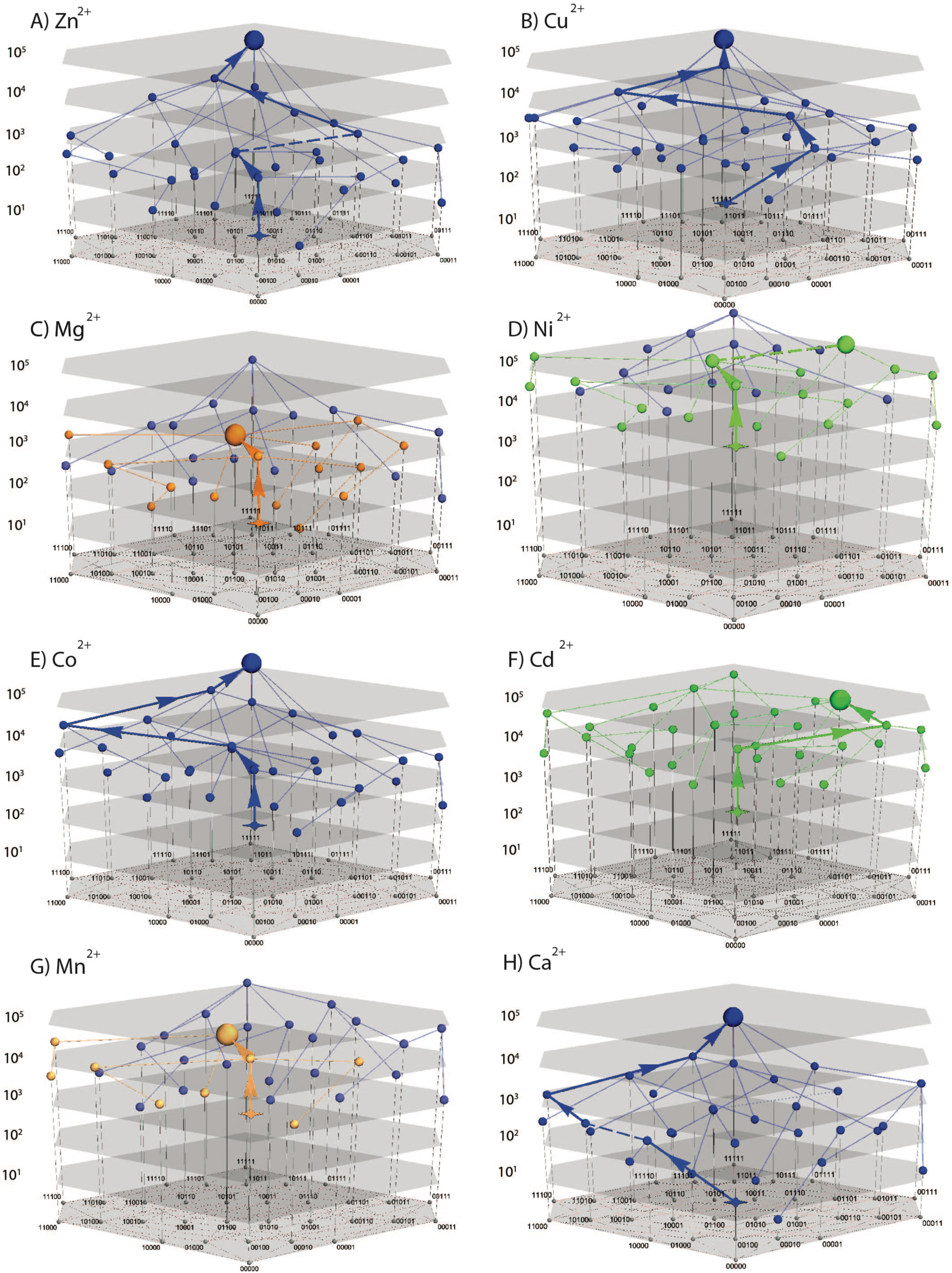
**A-H)** The adaptive landscape encompassing all 32 genotypes that define this evolutionary transition for all metal environments tested. Nodes reflect position in genotype space (mapped along bottom using binary encoding for all five substitions, 0 = ancestral state, 1 = derived state). Vertical height reflects catalytic activity (in cell-normalized nM/sec). Local and global optimal genotypes are highlighted with larger nodes while the ancestral genotype (DHCH) is highlighted by a star node. Dashed lines or arrows indicate transitions that are within the margin of error. Blue nodes and lines indicate those that reach the derived genotype (11111); red nodes and lines indicate those that reach the second most common optimal genotype (01100); green nodes and lines indicate those that reach the third most common optimal genotype (01101). Note that the second and third most common optima are connected by a single mutation and, for most metals, are relatively close in value, suggesting they might more accurately be described as a single “ridge” rather than two distinct peaks.

### Secondary environmental variation alters mutational effects and key epistatic interactions

Why do different metal environments produce unique adaptive landscapes and distinct evolutionary outcomes? In particular, what is the molecular basis underlying the unique topology of Cd^2+^ adaptive landscape? It is expected that evolutionary trajectories can be impacted by two non-additive phenomena: First, genotype-by-environment (*G×E*) interactions (which change in the effect of single point mutations in different environments)^32^ and second, genotype-by-genotype-by-environment (*G×G×E*) interactions (which imply that specific epistatic interactions vary depending on the environment).^33^ In order to quantify the impact of metal environment on each historical mutation, we first determined the average effect of each mutation across all possible genotypic backgrounds (see Methods).^34,35^ As we described previously, the effect of each mutation on MPH activity varies substantially depending on existence other mutations, suggesting extensive epistatic interactions among the five mutations.^23^ Different secondary environments resulted in qualitatively similar average single mutational effects, with each mutation usually either increasing (*l*72R, Δ193S, *h*258L, and *f*273L) or decreasing (*i*271T) enzyme activity (Figure 5). The magnitude of each mutation’s effect, however, varied depending on secondary environment. For example, *l*72R has a highly positive effect in Zn^2+^, Cu^2+^, Co^2+^ and Ca^2+^ environments, but only a marginal effect in Mg^2+^, Ni^2+^, Cd^2+^ environments, and a slightly negative effect in the Mn^2+^ environment, indicating that *G×E* interactions at least partially explain variation in the adaptive landscapes. Interestingly, however, the similar average effects of the five individual mutations in all metal environments, including Cd^2+^, suggest that *G×E* interactions alone are insufficient to explain the uniqueness of the Cd^2+^ adaptive landscape (Figure 4 and 5).

**Figure 5:**
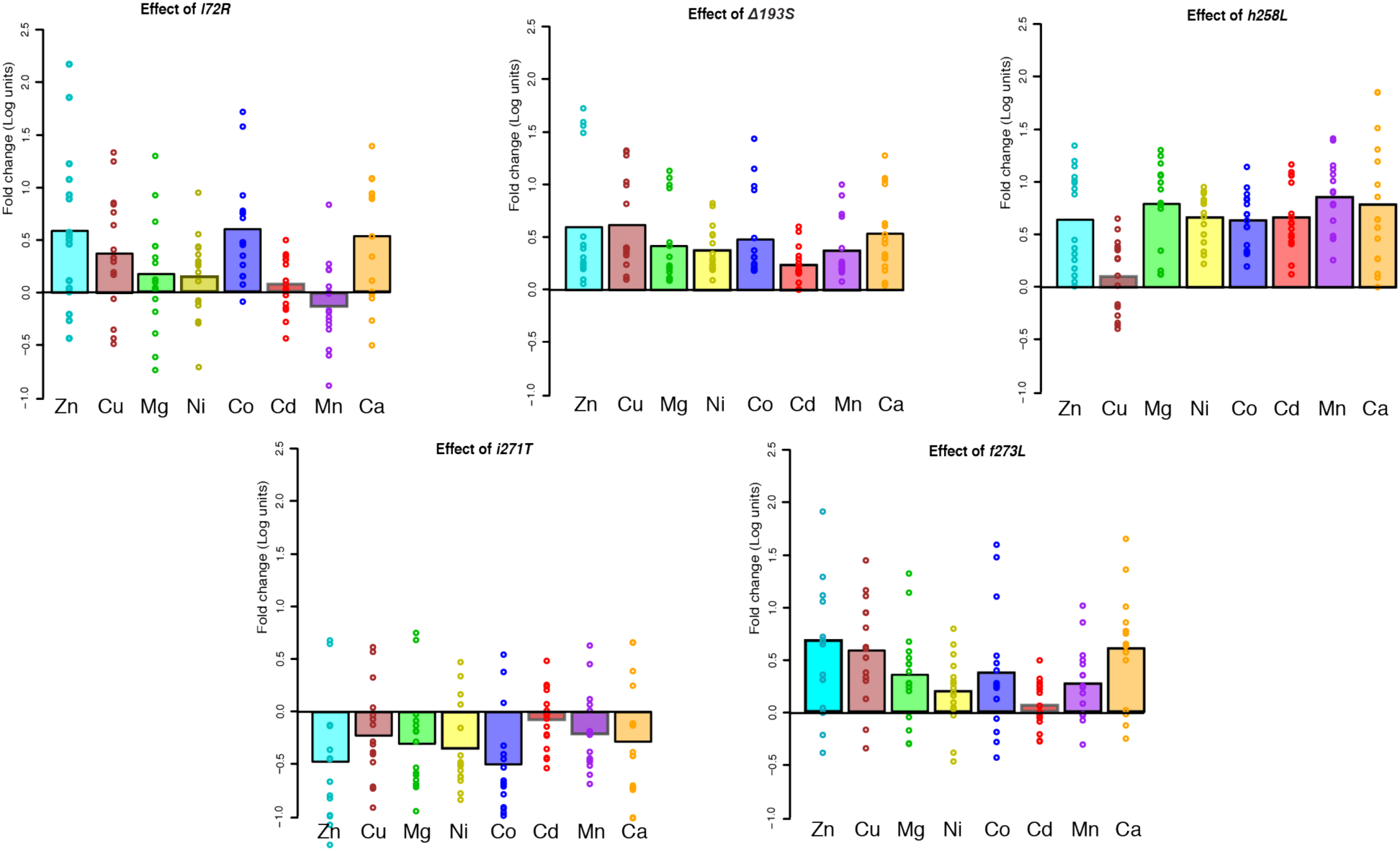
The average mutational effect of each of the five key substitutions for each metal environment (shown in solid bar). The effect of each mutation introduced in a specific genetic background is shown with dots.

Next, we examined the effect of each mutation when introduced into all 16 alternative genetic backgrounds (*i.e.*, its epistasis) and performed pairwise linear regression of those effects in different metal environments to assess how well correlated overall epistasis is between environments (equivalent to genotype-by-genotype-by-environment, or *G×G×E*, interactions: see **Methods**).^36^ Further, we constructed a more complex linear model to fit the adaptive landscape to calculate the degree and contribution of epistasis, including higher order epistasis, in each metal environment, and determine the impact of the secondary environment on epistasis. The contribution of epistasis is similar across the all metal environments: the first order effect of mutations explain 68%-78% of the overall variation in activity, while between 21%-31% is attributable to epistasis. However, first- and second-order epistasis (i.e. a model that includes average effects and pairwise epistatic interactions) explains between 87%-97% of the overall variation in activity, while higher order epistasis (3^rd^-5^th^ order) contributes only 0.5-10% collectively (Supplemental Table 2). When we examine the degree of second order epistasis, we found significant variation in both the magnitude and the sign (*i.e.*, switching from increasing to decreasing catalytic activity, or vice versa) of specific epistatic interactions across environments (Figure 6A). For example, the *h*258L*×i*271T interaction is highly synergistic in the Zn^2+^, Mg^2+^, Cu^2+^ and Ca^2+^ environments, but only marginal in the Mn^2+^, Ni^2+^ and Co^2+^ environments, and is highly antagonistic in the Cd^2+^ environment. Similarly, the *l*72R*×f*273L and *i*271T*×f*273L interactions are positive for all metal environments except Cd^2+^ (Figure 6A). A set of smaller individual effects can explain the difference in other metal environments. For example, smaller average effects of *l*72R, Δ193S and *f*273L (*G×E)* as well as the less synergistic *h*258Lx*i*271T and Δ193S-*f*273L interactions contribute to the difference in overall improvement by all five mutations in Zn^2+^ and Ni^2+^ environment (910-fold *vs*. 18-fold, Figure 2C).^11^ These GxE and GxGxE interactions help to explain the unique topology of each metal’s adaptive landscape, demonstrating that they can profoundly alter the evolutionary trajectories across it.

**Figure 6:**
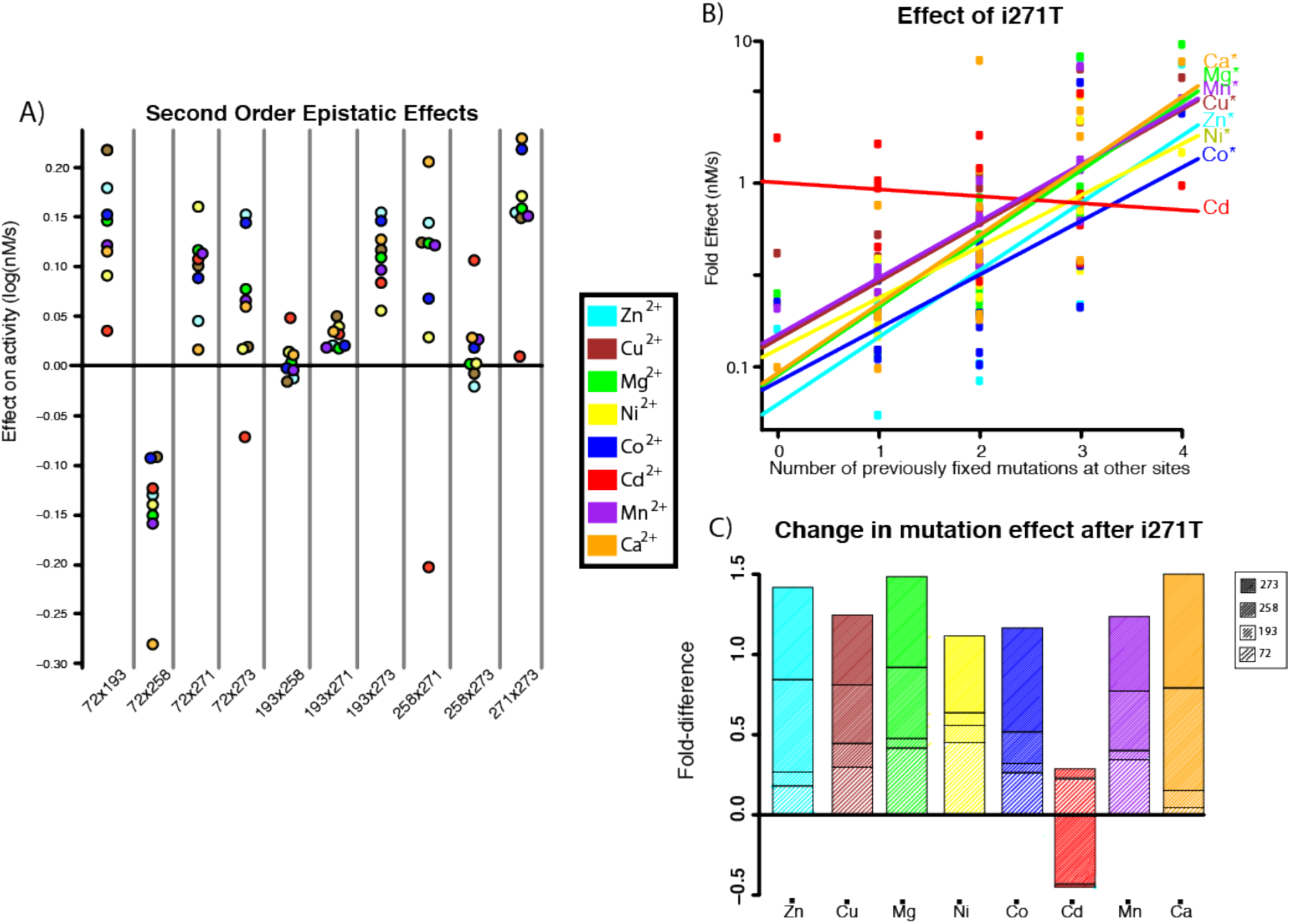
A) Shows the pairwise epistatic interaction effects for each metal environment. Cd^2+^ is a clear outlier for interactions between 258×271 (for which it exhibits the only negative interaction) and between 271×273 (for which it exhibits a uniquely marginal interaction while all other metal environments have strongly positive interactions). B) Shows the relationship between the effect of mutating position 271 and the number of previously fixed mutations at other sites for each metal. * denotes a statistically significant correlation. Note that all metal environments show a significant correlation except for Cd^2+^. C) Shows the impact of previous mutations at positions 72, 193, 258 and 273 on the effect of the mutation at position 271. Note that over 50% of the variation in the effect of the mutation at position 271 is determined by positive correlations with mutations at positions 258 and 273 for all metal environments except for Cd^2+^.

### Unique *G×G×E interactions lead to different evolutionary trajectories*

Last, we sought to further investigate the molecular basis underlying the differences in the evolutionary trajectories under different metal environments. As previously described, with the exception of Cd^2+^, all seven other metal environments have the highest activity across this set of sequence space at the fully derived MPH genotype. However, the topology of each landscape is still unique, as each metal environment results in a distinct evolutionary trajectory beginning from the fully ancestral genotype (Figure 7); while some can clearly evolve to the MPH genotype, others could potentially become stranded at a different genotype representing a local maximum instead (Figure 4C,D,F,G). We analyzed the mutational effects and epistatic interactions that were responsible for these different adaptive landscape topologies. We note several key *G×G×E* interactions that at least partially explain how these landscape differences emerged: *i*271T is of particular interest, as this mutation reduces activity when introduced into the ancestral genetic background in all metal environments except Cd^2+^, only becoming positive several other mutations have first arisen (*i*271T’s effect on activity is positively correlated with the number of mutations that were previously fixed for all environments except Cd^2+^ – Figure 6B). Furthermore, this pattern of *i*271T’s dependence on other mutations is driven by its interactions with *h*258L and *f*273L (Figure 6C). For example, epistasis does lead to an increase in the effect of *i*271T in the Mn^2+^, Mg^2+^ and Ni^2+^ environments as other mutations arise, however, these interactions are not strong enough to reverse the sign of *i*271T from negative to positive (Figure 3C), explaining why adaptive trajectories in those environments fail to reach the fully derived MPH genotype. Moreover, the two mutations, *l*72R and *f*258L, that initially cause the greatest increase in activity (Figure 3C) become negative if introduced after the other four mutations due to epistasis. For example, *l*72R-*h*258L exhibits strong antagonistic epistasis and the fixation of *h*258L leads to *l*72R decreasing enzyme activity in some environments, similarly preventing those trajectories from reaching the fully derived MPH genotype (Figure 6A, 7).

**Figure 7:**
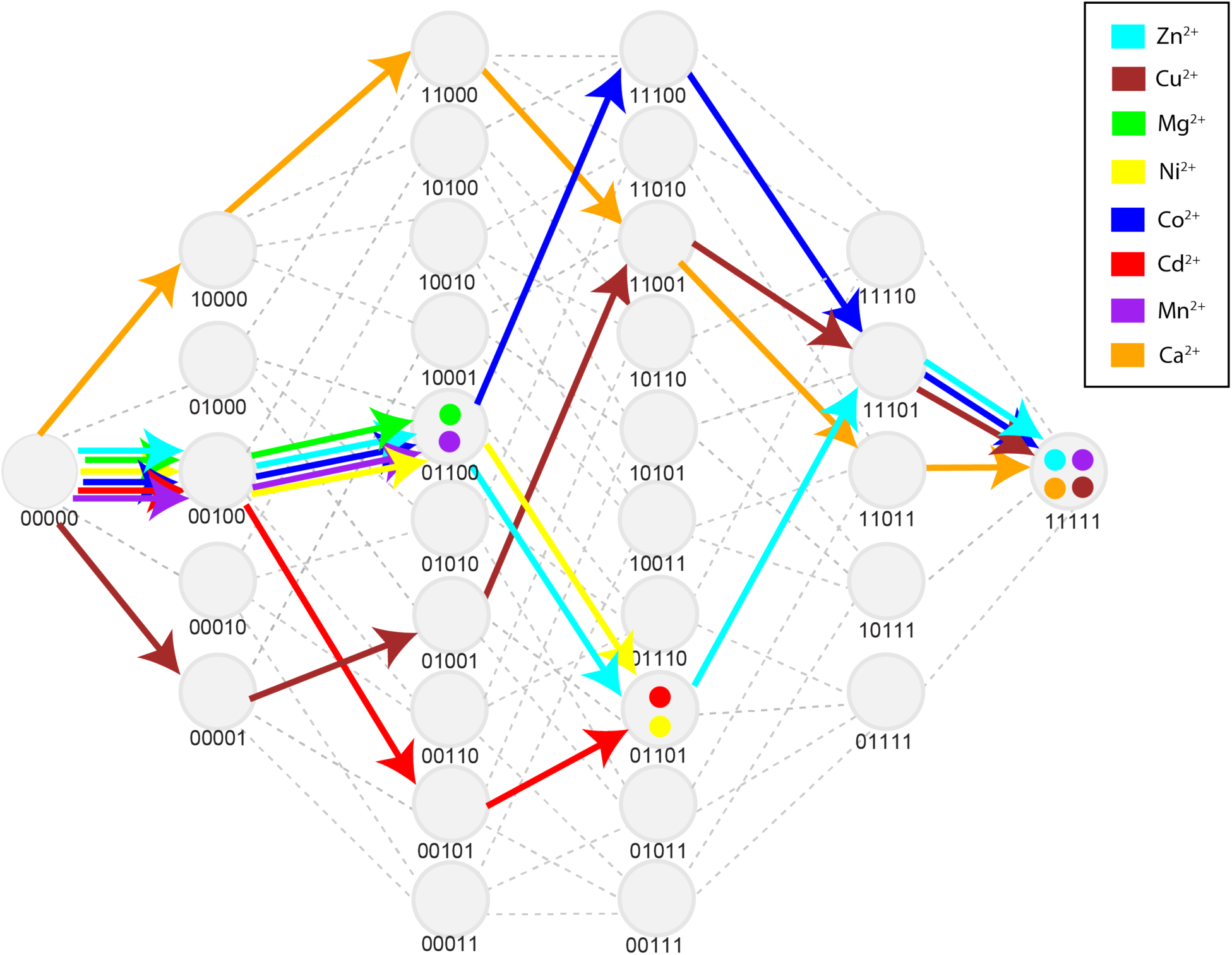
Displays the specific mutations as they are accumulated along the adaptive trajectory for each metal environment. Coloured dots indicate the corresponding metal environment’s “end point” for the trajectory beginning at the ancestral DHCH genotype. This corresponds to each landscape’s either global or a local optimum.

Taken together, the different topology of adaptive landscapes and the existence of some local optima are the result of several different *G×G×E* interactions. In one case, the degree of synergistic epistasis causes an initially negative mutation (*i*271T) to become positive, thus opening a newly accessible mutational pathway. In other cases, antagonistic epistasis causes initially positive mutations (*l*72R and *f*273L) to become negative, thereby restricting potential mutational pathways. This highlights the prominence of epistatic interactions in directing evolutionary trajectories, and demonstrates how even small shifts in the magnitude or sign of those interactions can result in different adaptive outcomes (Figures 4, 6–7).

## Discussion

Overall, this work demonstrates that the secondary environment of metal abundance can significantly alter evolutionary trajectories and outcomes in the case of adaptation to increased methyl-parathion hydrolysis from the DHCH ancestral enzyme tested here. In particular, our observations reveal several critical details about the effect of secondary environmental variation on both adaptive landscapes and the evolutionary trajectories that traverse them. First, alternative secondary environments can possess critical differences in both the quantitative measure of a protein’s function and on the direction and magnitude of mutational effects (*G×E* interactions). Second, and even more important from an evolutionary standpoint, is that secondary environmental variation can dramatically alter specific epistatic interactions, (*G×G×E* interaction) in some cases causing complete sign reversal between environments. Finally, the consequence of these changes on epistasis and the adaptive landscape lead to changes in the potential evolutionary outcome.^37,38^ In one case, it changes the genotype of the global optimum across this set of evolutionary sequence space, while in others it may instead reach a local optimum as the protein evolves across the adaptive landscape.

What could be the molecular basis for these unique epistatic interactions, adaptive landscapes, and evolutionary outcomes? Our previous work suggested that the distinct electrostatic properties of the metal ions, rather than any radical change in the active site, caused different activity profiles of the fully derived MPH enzyme by subtly altering substrate and transition state geometries.^25^ Moreover, we have also previously shown that these five historical mutations increase the methyl-parathion activity by repositioning the substrate through changing of the shape of the active site cavity.^23^ Thus, none of the genotypic and metal environmental changes drastically alter the mechanism of the MPH’s catalysis; instead, it is likely that each subtle change of the electrostatic and/or active site cavity acts in concert to fine-tune the alignment between the substrate and catalytic machinery. Consequently, changes in even small physical positions can impact key epistatic interactions, thereby altering the topology of the adaptive landscape and leading to different adaptive outcomes.

Our analyses have several shortcomings that are worth noting. First, the secondary environments that we tested, while based on environmentally realistic metal concentrations, are unrealistically simple – the true environmental variation likely spans many different combinations of divalent metal ions that could alter our conclusions about adaptive landscapes and adaptive outcomes,^37^ as well as other factors such as different pH, temperature, and salinity.^11^ Second, the Darwinian model of strong directional selection for a maximized catalytic function is applied only to the set of five mutations we identified as being responsible for MPH adaptation – in reality, the adaptive landscape would almost certainly have included many other potential mutations, and likely a myriad of alternative potential pathways.^39,40^ These shortcomings, however, are most likely to minimize the significance of our observations here, and exploring each of them more fully will likely reveal even greater sensitivity and variability in the process of adaptation, and the existence of even broader (and more variable) evolutionary watersheds defined by an even greater range of secondary environmental variation.^41^

How common are such shifting evolutionary watersheds likely to be in other systems? At this point, we can only speculate, as data on many more systems must first be collected and analyzed in order to definitively resolve this question. In the case of the DHCH-to-MPH evolutionary transition, the secondary environment of metal abundance is directly linked to replacement of the cofactor in the active site.^24,25^ A large fraction of known enzymes are metalloenzymes, many of which have been shown to bind promiscuously to different metal ions that alter their activity profiles.^42^ Additionally, many enzymes utilize other cofactors and bind to different types of cofactors.^43–45^ Moreover, other environmental factors such as temperature, pH, redox potential, salinity, and expression of other proteins such as chaperones, can impact enzyme function and expression, and thus the effects of specific mutations on their function.^46–51^ Therefore, the secondary environment could easily play a similarly significant role in many other cases of molecular adaptation. We propose that further studies examining these phenomena should emulate the approach we have used here by examining genetic and environmental effects in concert in order to assess the different shapes that an adaptive landscape may take. Combining the careful construction of statistical linear models and detailed evolutionary pathway analyses under reasonable models of evolution can allow us to more clearly assess the impact that *G×E* and *G×G×E* interactions have on evolving proteins. By undertaking this task, we can characterize not only the adaptive landscapes defined by key genetic changes, but we can assess their sensitivity to secondary environmental variation, thereby etching out the contours of the evolutionary watersheds within them.^52^

When there is significant secondary environmental variation and prominent mutational epistasis, evolutionary watersheds can shift, becoming contingent on the conditions in which evolution occurs. Thus, it is critical that we carefully consider the secondary environment as well as the genotypic background in our efforts to predict, design, and understand the evolution of new biological molecules. Adaptation reflects the conditions in which it occurs: its outcome, like water draining into the ocean, depends both on where it begins and on the landscape across which it travels.

## Methods

### Enzyme cloning and kinetic measurements

Enzyme genotypes were mutated and cloned into a pET27(b) vector (Novagen) containing a N-terminal Strep-tag II sequence (MASWSHPQFEKGAG) using the *Nco* I and *Hind* III restriction enzymes (Thermo Scientific), as described previously.^23^ To test the lysate activities of variants, *E. coli* Bl21 (DE3) transformed with plasmids for each of the 32 MPH variants were grown in triplicates in a 96-deep well plate containing 200 µL of LB media supplemented with 40 µg/mL kanamycin at 30°C, 900 × rpm overnight. On the following day, a second 96-deep well plate containing 400 µL of LB media supplemented, 50 µg/mL kanamycin, and 100 µM of one of the eight metal ions were inoculated with 20 µL of the aforementioned overnight culture and incubated at 30°C, 900 × rpm for 3 hours. Protein expression was induced by adding IPTG to a final concentration of 1 mM and further incubation at 30°C for 3 hours. Cells were harvested by centrifugation at 3,220 × g for 10 minutes and pellets were frozen −80°C for at least 30 minutes. To lyse the cells, the cell pellets were resuspended in 200 µL of lysis buffer consisting of 50 mM Tris-HCl pH 7.5, 100 mM NaCl, 200 µM of the same metal ion that was supplied in the LB, 0.1% Triton X-100, 100 µg/ml lysozyme and 1 U/ml of benzonase, and incubated at room temperature with shaking at 1200 × rpm for 1 hour. The cell lysates were clarified by centrifugation at 3,220 × g for 20 minutes at 4°C. Clarified lysates were diluted 5-fold and measured against a single substrate concentration of 400 µM for the methyl-parathion substrate to obtain linear initial rates. The lysate activity is given as the rate of substrate hydrolysis in µM/sec/OD, which is calculated from the molar extinction coefficient of the *p*-nitrophenol leaving group (18,300 M^−1^cm^−1^) and normalized to the OD of the cell cultures.

### Linear modeling genetic and environmental effects

#### Definition of genetic and environmental encoding system

To quantify the genetic and environmental determinants of enzyme activity, we used an approach similar to that previously developed.^34,35^ We constructed regression models that explain lysate activity (*L.A.*) as a function of the genetic states at the five variable amino acid residues in the protein.

The genetic variation in the protein was defined in the linear models using one-dimensional variables for the mutations; residues 72, 193, 258, 271, and 273 are described by single-dimensional vectors a, b, c, d, and e, respectively, with the ancestral state defined as −1 and the derived state defined as +1 These variables make the y-intercept of the linear model equal to the mean activity across all experimental measurements ^35^; therefore, all genetic effects are expressed relative to the mean (Supplemental Table 1).

#### First-order linear models

We constructed our first-order model by regressing the lysate activity of each genotype on dependent variables that reflect the individual first-order identities at each genetic position. For example, the linear model for position 72 is expressed as:

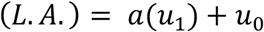

where *a* is the effect coefficient of moving +1 in that dimension, *u1* is the coordinate representing the genotype (i.e. −1 for ancestral leucine, +1 for derived arginine), and *u*;_0_ is the y-intercept for the model (equal to the mean across the data). The linear coefficients for each model were computed using ordinary least squares (OLS) regression with the open-source statistical package R (http://www.r-project.org/). The coefficient *a* indicates the deviation of the derived genetic state from the mean, while *–a* gives the deviation of the ancestral genetic state from the mean.

To determine how well all five first-order effects of mutations in the protein predict variation in *L.A.*, we constructed the following linear model that included all first-order protein coefficients:

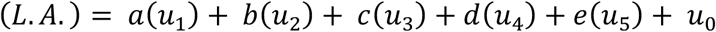

where *u*_2_, *u*_3_, *u*_4_, and *u*_5_ are the coordinates representing the genotype for positions 193, 258, 271, and 273, respectively. We then computed the R^2^ for this first-order model.

The first-order models for the effect of each environmental factor (i.e. which metal ion was present in the lysate) was modeled using expanded variable space applied along the lines described previously.^34,35^ For this, each metal variable is assigned a unique set of coordinates in 7-dimensional space according to the relevant Hadamard matrix, and those variables were then used to perform a minimal-variable linear regression that is similarly centred to the mean across all the data:

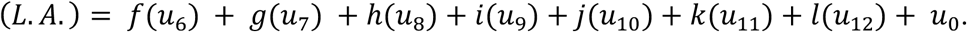

Where *u*_6_, *u*_7_, *u*_8_, *u*_9_, *u*_10_, *u*_11_, and *u*_12_ are the coordinates representing the metal contained in the lysate (full datasets and computational scripts available on Github: https://github.com/danderson8/Evolutionary-Watersheds). The magnitude of the effect of each metal on lysate activity was determined by computing the sum of the modeled coefficients for its defined coordinates.

#### Linear models with second-order genetic epistasis and G x E interactions

To identify cases of second-order epistatic interactions and genotype by environment (G x E) interactions, we individually introduced every possible interaction term for every two-way combination of genotypes at the variable sites in the protein or the metal environment. These interaction variables were constructed as previously described.^34,35^ Each interaction is described by a new linear vector, the value for which is determined by taking the outer product between the two first-order linear vectors. For example, the interaction between site 72 and 256 of the protein will be equal to (*a*) ⊗ (*c*) = (*ac*).

Where *u*_13_ is equal to *u*_1_*u*_3_, etc. The second-order interaction effects are equal to the deviation from the additive effect modeled by each genetic state individually across other genetic backgrounds, and is defined herein as the “marginal” effect (i.e. added on to the “average” effects computed in the first-order model). Interactions between each mutation and the metal environment were modeled analogously. For example, the interaction between site 72 in the protein and the metal environment is constructed by: (u_1_) ⊗ (u_6_, u_7_, u_8_, u_9_, u_10_, u_11_, u_12_) = (u_1_u_6_, u_1_u_7_, u_1_u_8_, u_1_u_9_, u_1_u_10_, u_1_u_11_, u_1_u_12_).

One advantage of this method of encoding the genetic data is that the first-order model is nested within the second-order model. This allowed us to assess whether addition of the second-order model terms significantly improved the fit by comparing the improvement in the adjusted R^2^ as well as the improvement in the likelihood ratio test relative to the simpler first-order model. The effect of each second-order interaction (i.e. the epistasis and/or GxE interactions that should be added to the sum of the additive lower-order effects) can be solved from these coefficients.

### Evolutionary pathway determination

To model the evolutionary pathway under a model of strong direction selection (i.e. classic Darwinian adaptation), we developed our own in-house script that calculates the most likely evolutionary pathway under a model of Darwinian selection (available here: https://github.com/danderson8/Evolutionary-Watersheds). Briefly, each genotype’s triplicate measurements are considered, and for each genotype the difference in the average activity between it and its single-mutational neighbours (including reversals) is considered for the five evolutionarily relevant changes (specified in the main text). Whichever neighbouring genotype provides the greatest improvement in MPH activity (as long as the change is not negative) is selected as the most likely evolutionary “step”. An assessment of the confidence in each step is then made by comparing the span of replicate measurements for each of the two genotypes: if they overlap then the mutational step is considered “ambiguous”, and it is represented in the trajectory diagram as a dotted or dashed line (see Figure 4). The “new” genotype is then analyzed in the same way. The process is repeated with each subsequent mutational “step” until an optimal genotype is reached, for which all the neighbouring genotypes are significantly (i.e. non-overlapping) lower in their MPH activity measured, which meets the conditions of being either a local or global optimum across the landscape.

---

**Supplemental Figure 1:**
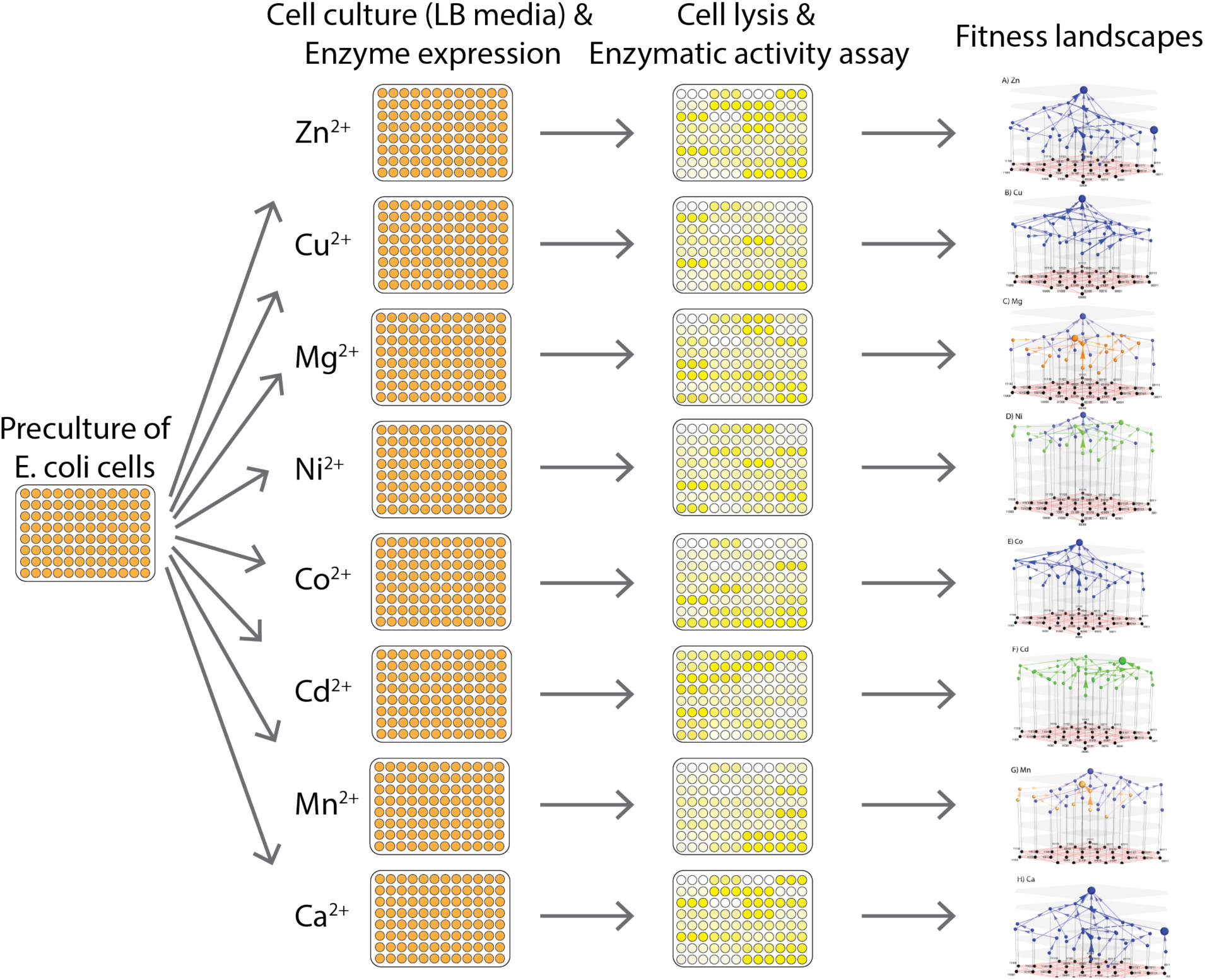
Outline of the experimental and analytical scheme.

**Supplemental Table 1:** All mutational effects as determined by linear models (see **Methods**) across all metal environments tested. Epistatic interactions were determined up to full fifth-order interactions.

| Terms | Zn | Cu | Mg | Ni | Co | Cd | Mn | Ca |
| --- | --- | --- | --- | --- | --- | --- | --- | --- |
| pos72 | 0.295 | 0.188 | 0.083 | 0.070 | 0.297 | 0.035 | -0.063 | 0.264 |
| pos193 | 0.301 | 0.311 | 0.211 | 0.190 | 0.242 | 0.120 | 0.188 | 0.269 |
| pos258 | 0.321 | 0.051 | 0.397 | 0.333 | 0.318 | 0.333 | 0.429 | 0.394 |
| pos271 | -0.236 | -0.111 | -0.151 | -0.172 | -0.248 | -0.036 | -0.103 | -0.140 |
| pos273 | 0.338 | 0.291 | 0.175 | 0.097 | 0.185 | 0.031 | 0.133 | 0.300 |
| pos72pos193 | 0.179 | 0.161 | 0.132 | 0.064 | 0.113 | 0.018 | 0.094 | 0.090 |
| pos72pos258 | -0.129 | -0.068 | -0.135 | -0.098 | -0.069 | -0.065 | -0.121 | -0.218 |
| pos72pos271 | 0.045 | 0.074 | 0.104 | 0.112 | 0.066 | 0.056 | 0.086 | 0.011 |
| pos72pos273 | 0.151 | 0.014 | 0.069 | 0.011 | 0.108 | -0.037 | 0.050 | 0.046 |
| pos193pos258 | -0.012 | -0.012 | 0.003 | 0.008 | -0.001 | 0.025 | -0.003 | 0.008 |
| pos193pos271 | 0.021 | 0.037 | 0.015 | 0.027 | 0.014 | 0.017 | 0.014 | 0.027 |
| pos193pos273 | 0.154 | 0.086 | 0.098 | 0.038 | 0.110 | 0.044 | 0.074 | 0.098 |
| pos258pos271 | 0.144 | 0.092 | 0.111 | 0.020 | 0.049 | -0.107 | 0.093 | 0.160 |
| pos258pos273 | -0.020 | -0.006 | -0.001 | 0.000 | 0.014 | 0.056 | 0.020 | -0.021 |
| pos271pos273 | 0.155 | 0.109 | 0.142 | 0.120 | 0.162 | -0.004 | 0.116 | 0.178 |
| pos72pos193pos258 | -0.005 | 0.037 | 0.007 | 0.010 | 0.001 | 0.027 | -0.013 | 0.038 |
| pos72pos193pos271 | 0.030 | 0.045 | 0.029 | 0.018 | 0.028 | 0.026 | 0.002 | 0.030 |
| pos72pos193pos273 | 0.167 | 0.064 | 0.084 | 0.044 | 0.103 | 0.016 | 0.062 | 0.094 |
| pos72pos258pos271 | 0.006 | -0.002 | 0.058 | 0.033 | 0.042 | 0.022 | 0.008 | -0.044 |
| pos72pos258pos273 | 0.083 | 0.104 | 0.014 | 0.019 | 0.041 | 0.041 | -0.007 | 0.059 |
| pos72pos271pos273 | 0.142 | 0.124 | 0.060 | 0.073 | 0.098 | 0.045 | 0.031 | 0.054 |
| pos193pos258pos271 | -0.004 | -0.007 | -0.006 | -0.009 | -0.005 | -0.015 | -0.020 | -0.032 |
| pos193pos258pos273 | 0.017 | 0.014 | -0.003 | -0.011 | -0.019 | 0.026 | -0.001 | -0.019 |
| pos193pos271pos273 | -0.025 | -0.028 | -0.010 | -0.033 | -0.013 | 0.011 | -0.001 | -0.015 |
| pos258pos271pos273 | 0.072 | 0.041 | 0.052 | -0.010 | 0.058 | 0.016 | 0.041 | 0.049 |
| pos72pos193pos258pos271 | -0.010 | -0.001 | -0.017 | -0.024 | -0.019 | -0.024 | -0.017 | 0.049 |
| pos72pos193pos258pos273 | -0.008 | -0.007 | -0.012 | -0.005 | -0.012 | 0.014 | 0.007 | -0.005 |
| pos72pos193pos271pos273 | -0.001 | -0.025 | -0.004 | -0.017 | 0.012 | 0.003 | 0.000 | -0.005 |
| pos72pos258pos271pos273 | 0.005 | -0.028 | -0.016 | -0.014 | 0.006 | -0.012 | 0.002 | -0.004 |
| pos193pos258pos271pos273 | 0.002 | -0.019 | 0.004 | -0.019 | -0.032 | 0.005 | -0.008 | 0.002 |
| pos72pos193pos258pos271pos273 | -0.022 | -0.014 | 0.006 | -0.020 | -0.026 | -0.012 | -0.014 | 0.012 |

**Supplemental Table 2:** R^2^ values for all linear models tested. These reflect the proportion of overall variation (including experimental error) that is explained by a linear relationship with the specified mutation or the respective model (non-epistatic = first order, pairwise epistasis only = second order, higher-level epistasis = third through fifth orders). For second-fifth order models the R^2^ specified is the increased R^2^ that results from fitting the more complex model as compared to the next highest-order model.^34^

| Terms | Zn | Cu | Mg | Ni | Co | Cd | Mn | Ca |
| --- | --- | --- | --- | --- | --- | --- | --- | --- |
| pos72 | 0.13 | 0.10 | 0.018 | 0.020 | 0.20 | 0.035 | 0.012 | 0.12 |
| pos193 | 0.14 | 0.29 | 0.12 | 0.15 | 0.13 | 0.120 | 0.11 | 0.13 |
| pos258 | 0.16 | 0.008 | 0.42 | 0.45 | 0.22 | 0.333 | 0.57 | 0.28 |
| pos271 | 0.084 | 0.037 | 0.060 | 0.12 | 0.14 | -0.036 | 0.033 | 0.035 |
| pos273 | 0.17 | 0.25 | 0.082 | 0.038 | 0.076 | 0.031 | 0.055 | 0.16 |
| First Order Model | 0.68 | 0.68 | 0.70 | 0.77 | 0.76 | 0.018 | 0.78 | 0.72 |
| Second Order Model | 0.21 | 0.19 | 0.25 | 0.18 | 0.17 | -0.065 | 0.19 | 0.22 |
| Third Order Model | 0.094 | 0.11 | 0.048 | 0.042 | 0.06 | 0.056 | 0.022 | 0.04 |
| Fourth Order Model | 0.0003 | 0.005 | 0.002 | 0.006 | 0.004 | -0.037 | 0.001 | 0.004 |
| Fifth Order Model | 0.0007 | 0.0006 | 0.0001 | 0.002 | 0.001 | 0.025 | 0.0006 | 0.0003 |

